# Auditory binaural and spatial hearing impairments in a Fragile X Syndrome mouse model

**DOI:** 10.1101/648717

**Authors:** Elizabeth A. McCullagh, Shani Poleg, Nathaniel T. Greene, Molly M. Huntsman, Daniel J. Tollin, Achim Klug

## Abstract

The auditory brainstem compares sound-evoked excitation and inhibition from both ears to compute sound source location. Although alterations to the anatomy and physiology of the auditory brainstem have been demonstrated in Fragile X Syndrome (FXS) it is not known whether these changes cause sound localization impairments in FXS. To test the hypothesis that FXS-related alterations to brainstem circuits impair spatial hearing abilities, a reflexive prepulse inhibition (PPI) task, with sound source location as the prepulse stimulus, was used to show that *Fmr1* knockout mice have decreased inhibition of their startle responses. Specifically, *Fmr1* mice show decreased PPI compared to wildtype during gap detection, changes in sound source location, and spatial release from masking with no alteration to their overall startle thresholds compared to wildtype. Lastly, *Fmr1* mice have increased latency to respond in these tasks suggesting additional impairments in the pathway responsible for reacting to a startling sound.

## Introduction

Fragile X Syndrome (FXS) is a neurodevelopmental disorder and the leading monogenetic cause of autism and mental retardation (Penagarikano et al., 2007). FXS is caused by a mutation in the gene *Fmr1* that encodes Fragile X Mental Retardation Protein (FMRP). One of the hallmark symptoms of FXS, among many other cognitive symptoms, is increased auditory hypersensitivity (reviewed in Rotschafer and Razak, 2014). An imbalance of neural excitation/inhibition (E/I) is thought to underlie many pathologies in FXS (Contractor et al., 2015) including those leading to auditory symptomology (Keine et al., 2016).

Recent work has shown that these E/I imbalances in FXS extend to the auditory brainstem circuits responsible for sound localization, as well as downstream cortical areas of the auditory system (Garcia-Pino et al., 2017; McCullagh et al., 2017; Rotschafer et al., 2015; Rotschafer and Cramer, 2017). FMRP is highly expressed in the auditory brainstem (Ruby et al., 2015; Wang et al., 2014; Zorio et al., 2017) leading to changes in potassium channel distribution (Brown et al., 2010; Strumbos et al., 2010) that underlie changes in synaptic function *in vitro* (Curry et al., 2018; El-Hassar et al., 2019; Garcia-Pino et al., 2017; Lu, 2019; Wang et al., 2015). In addition, studies have shown alterations to the auditory brainstem response (ABR), an *in vivo* measure of auditory brainstem activity, in *Fmr1* mice that are potentially caused by these underlying changes to E/I balance and physiological activity (El-Hassar et al., 2019; Rotschafer et al., 2015). Despite these clear and substantial alterations to the auditory brainstem, it has never been clearly shown that mice or humans with a mutation in *Fmr1* have impairments in their ability to localize sound.

Binaural hearing and spatial acuity are essential for communication in complex noisy acoustic environments, known as the “cocktail party problem” (Cherry, 1953). The separation of spatial channels in these complex noisy acoustic environments is dependent on an intricate E/I balance that starts in the auditory brainstem (Keine et al., 2016). Impairments in the E/I balance of the auditory brainstem sound localization pathway therefore are expected to lead to impaired communication ability in complex noisy acoustic environments. As a signal, or perhaps a speaker of interest, moves further in space from a distracting background noise, it becomes easier to discriminate from the noise, this effect is termed spatial release from masking (SRM)(reviewed in Feng and Ratnam, 2000).

This study aims to determine if mice with a mutation in the *Fmr1* gene have a functional deficit in sound localization. Sound source location discrimination was assessed using a reflexive prepulse inhibition (PPI) paradigm, where a change in the sound source location served as the prepulse, a method described previously for mice (Allen and Ison, 2010) and guinea pigs (Greene et al., 2018). The acoustic startle response is a reflexive whole-body response elicited by a very brief, but loud impulse noise. PPI consists of modification of the acoustic startle response by pairing the startle eliciting stimulus with a preceding auditory stimulus, the prepulse, that inhibits (i.e. attenuates) the startle response by providing a cue to the impending startle (Young and Fechter, 1983). PPI is a useful tool for measuring deficits in cognitive disorders such as autism, FXS, or schizophrenia since it is reflexive and independent of cognitive ability (Young and Fechter, 1983). It has been shown previously, with conflicting results, that mice and humans with *Fmr1* mutations have impaired PPI (Chen and Toth, 2001; Frankland et al., 2004; Hessl et al., 2009; Nielsen et al., 2002; Thomas et al., 2012; Veeraragavan et al., 2012) suggesting altered sensorimotor gating. We use a similar PPI paradigm, but with cues to specifically target the auditory brainstem (gap in sound, speaker swaps and spatial release from masking), to test the hypothesis that behavioral impairments resulting from altered signaling in the brainstem sound localization circuits (Allen and Ison, 2010). We show, for the first time, changes in *Fmr1* mice spatial acuity, quantified as reduced PPI and increased response latencies to startling sounds.

## Results

The results are based on experiments conducted in both male (N = 35) and female (N = 16) wildtype (B6) and *Fmr1* knock out (ko) mice. No significant differences were observed between sexes, and data for males and females were combined. Each animal was weighed after completion of data collection. Animal weights varied between 13-37 g, and generally increased with age. On average, *Fmr1* mice tended to weigh less (23.5 g) than wildtype animals (25.8 g) (p < 0.05).

### Startle threshold

Animals were initially tested to determine their response threshold to acoustic startle stimuli. This was done both to characterize the responses of *Fmr1* mice and ensure that all animals had a robust startle response with increasing intensity of sound. Startle amplitude is reported in units of arbitrary volts (output of the accelerometer) that are an uncalibrated measure of acceleration. Startle sounds were 20 ms in duration and varied in intensity between 60 and 120dB SPL were presented randomly, in 10dB SPL steps, in the presence of continuous 70dB SPL broadband noise (presented from the speaker directly in front of the animal) (Figure 1A). All animals were tested with intensities ranging from 80-120 dB SPL. Two additional levels (60 and 70 dB SPL) were tested in a subset of animals (6 B6, 8 *Fmr1*) to ensure that animals are not startled at lower intensity sounds. There was no difference between genotypes for either startle amplitude (Figure 1B, p=0.4074) or latency (Figure 1D, p=0.8331). Startle responses increased and latency decreased with increasing stimulus level in both genotypes, indicating that animals had no trouble detecting the startle stimulus and had a robust startle response. The magnitude of the startle responses for both B6 and *Fmr1* animals plateaued (threshold) around 100dB SPL and therefore the startle stimulus was set at 10 dB above this threshold (110dB SPL) for the remainder of the experiments.

**Fig 1.**
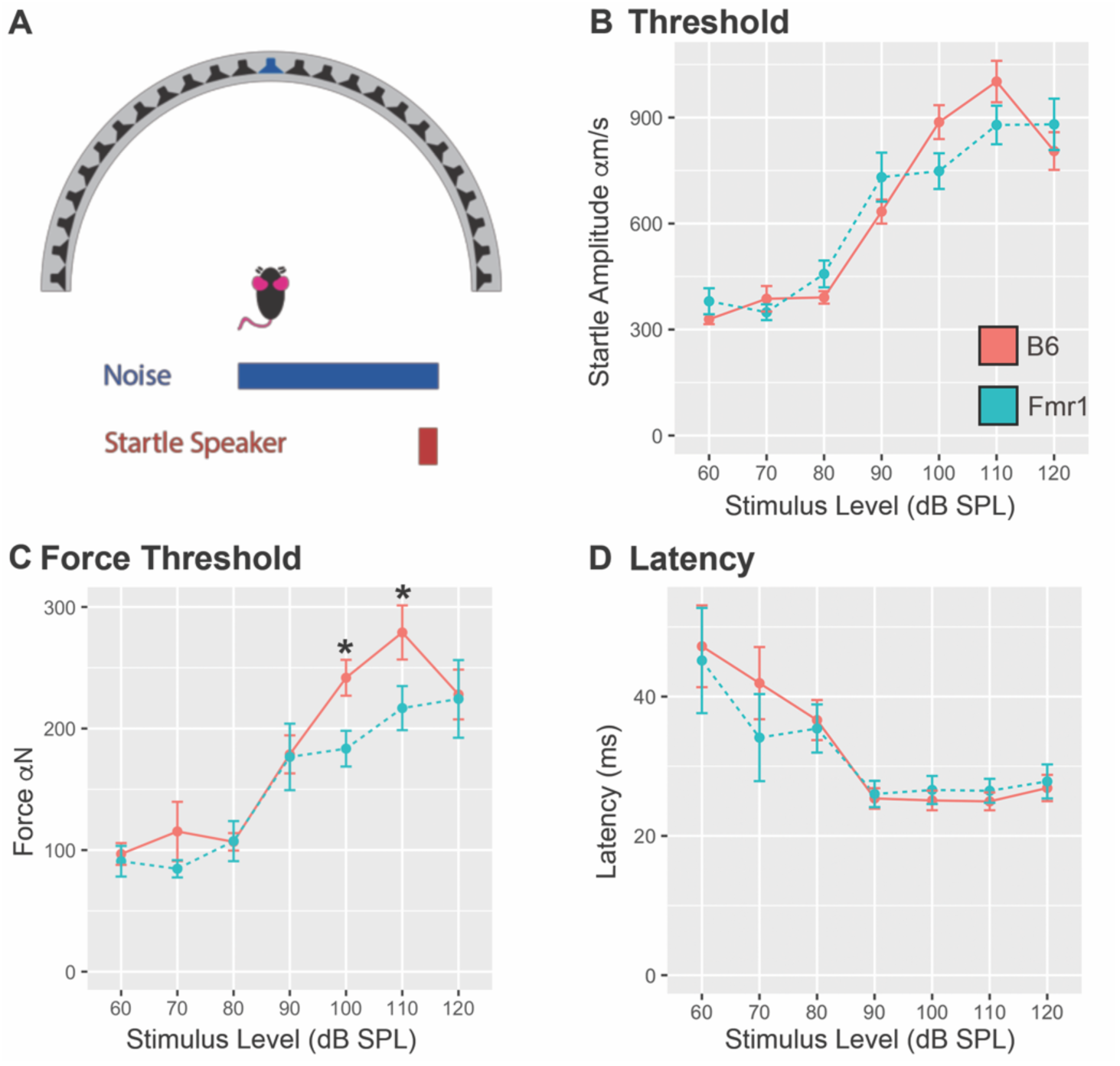
No difference in startle amplitude or latency between B6 and *Fmr1* mice, decreased force in *Fmr1* mice. A, illustration of the experimental setup. Animals were placed in the center of the speaker array in the presence of 70dB SPL broadband noise playing from the central speaker at 0° (blue speaker). Startle eliciting stimuli were presented by an additional speaker placed directly over the mouse’s head (not shown). B, the animal’s startle amplitude increased with stimulus level (dB SPL) for both B6 (red) and *Fmr1* mice (teal). A total of 22 B6 and 14 *Fmr1* mice were tested in this task, with a subset (6 B6 and 8 *Fmr1*) tested below 80 dB SPL. C, Force of response to the startle stimuli was smaller for *Fmr1* mice compared to B6. D, latency to respond (in ms) to the startle stimulus decreased with increasing stimulus intensity (dB SPL) for both B6 and *Fmr1* mice. Data shown are mean ± SEM for genotype. * p<0.05

When individual animal weights were used to calculate the force of each startle response, significant differences were observed between genotypes at 100 (p=0.0321) and 110 (p = 0.0224) dB SPL with *Fmr1* mice showing reduced startle compared to wildtype (Figure 1C). This could be due to a reduced muscle tone in *Fmr1* animals or some other factor, neither of which are explored further in this study. To account for differences in animal weight and reduced startle force we normalized responses are normalized to the baseline startle amplitude when calculating PPI (see methods).

### Gap Detection

Recent data has shown that impairments in gap detection may be caused by underlying changes to excitation-inhibition balance (E/I) in the inferior colliculus (IC) (Sturm et al., 2017), suggesting that gap detection may be used to probe the E/I balance in the auditory system. This E/I balance is known to be altered in FXS (Garcia-Pino et al., 2017; McCullagh et al., 2017; Rotschafer et al., 2015). We tested 42 animals (20 B6 and 22 *Fmr1*) in a gap detection paradigm with a 20 ms quiet gap in broad band noise (prepulse) followed by a startle eliciting stimulus at varying inter-stimulus interval (ISI) times (1-240 ms) between the prepulse and startle eliciting stimuli (Figure 2A). A subset of animals were only tested at 10-240 ms ISIs (8 B6, 8 *Fmr1*). There was no main effect of genotype (p=0.2011) at all levels of ISI, however there were significant differences between B6 and *Fmr1* animals at 10 ms (p<0.05) and 20 ms (p<0.01) ISIs (Figure 2B). In addition, *Fmr1* mice were slower to startle at all ISIs (as indicated by increased startle latency), except 10 ms and 20 ms, compared to B6 mice (Figure 2C). Both genotypes did not detect gaps less than 1 ms and *Fmr1* animals did not detect gaps of 2 ms, as indicated by PPI not significantly different than zero. These data suggest that not only do *Fmr1* mice have decreased PPI compared to wildtype at optimal ISIs but are also consistently slow to startle in most conditions, where in contrast, B6 mice show modulations to latency based on ISI.

**Fig 2.**
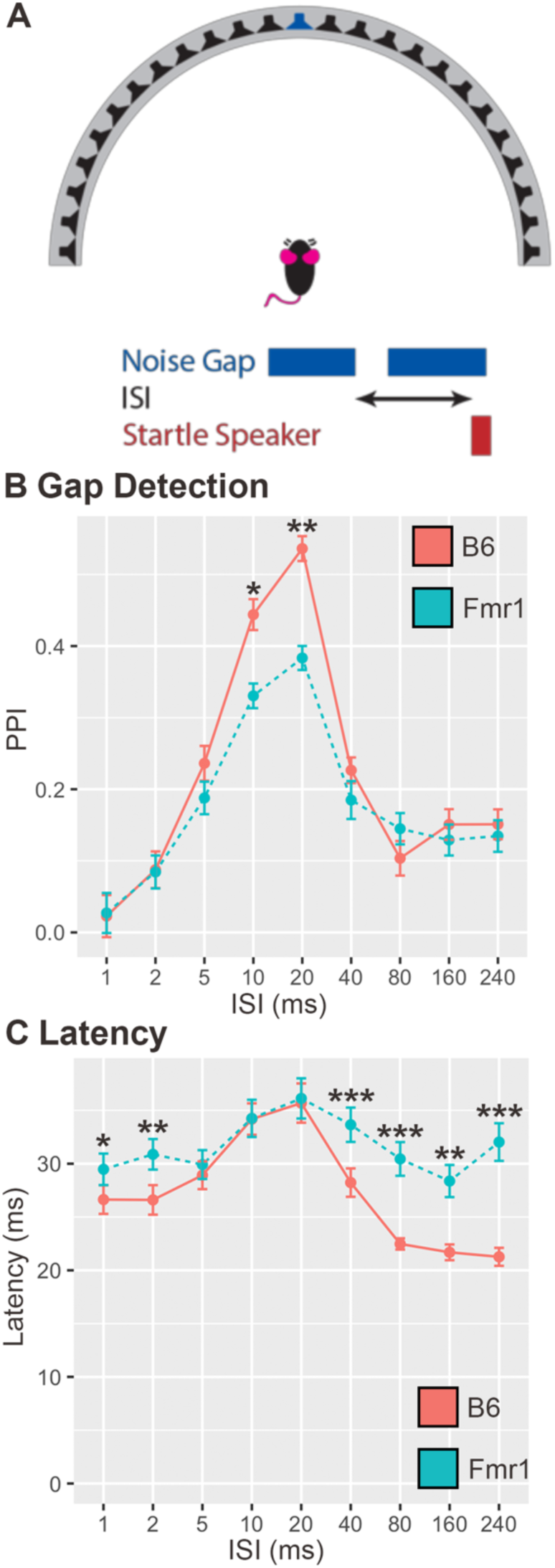
*Fmr1* mice have decreased PPI and increased latency to startle during a gap detection task. 42 mice (20 B6 and 22 *Fmr1*) were tested in a gap detection task, where a 20 ms quiet gap was introduced into 70dB SPL broadband noise (blue speaker at the center of the array), which was presented at varying ISIs before the startle eliciting stimulus presentation (A). *Fmr1* mice (teal) showed less PPI at 10 ms and 20 ms ISI compared to B6 (red; B). *Fmr1* mice startle responses latencies were longer to 1, 2, 5, 40, 80, 160 and 240 ms ISIs compared to wildtype (C). Data are shown as mean ± SEM. * p<0.05, ** p<0.01, *** p<0.001.

### Varying ISI with 90° speaker swap

The gap detection test suggests that *Fmr1* mice demonstrated deficits in temporal auditory processing, next we wanted to determine if *Fmr1* mice also have deficits in spatial auditory processing. The first step to determining whether *Fmr1* mice have spatial hearing deficits is to establish the optimal ISI for detection of a spatial speaker swap. In this task, the prepulse was a speaker swap of broadband noise from one speaker 45*°* to the right of the animal to the symmetrical speaker 45*°* to the left of the animal (90*°* total angle), with varying ISIs between the prepulse and startle eliciting stimuli (Figure 3A). There was no main effect of ISI on genotype (p=0.1068). However, *Fmr1* mice had reduced PPI of their startle compared to B6 mice at 20 ms and 30 ms (p<0.05) ISI after the 90*°* speaker swap (12 B6, 14 *Fmr1* mice, Figure 3B). In addition, *Fmr1* mice had increased latency to startle compared to B6 at all ISIs except 1 ms and 2 ms (Figure 3C). These data suggest that at ISIs that elicited some of the highest PPI for B6 animals (also similar ISIs that showed a deficit in the gap detection test), *Fmr1* mice showed reduced PPI compared to wildtype. Both genotypes did not detect, as indicated by a PPI not significantly different than zero, ISIs less than 5 ms. *Fmr1* animals did not detect the speaker swap (PPI not significantly different from zero) for ISIs of 10, 20, 30, 40, and 300 ms. In addition, *Fmr1* mice showed increased latencies to startle at ISIs that elicited a PPI above zero, suggesting that addition of a detectable prepulse actually slowed responses in comparison to B6 mice. Based on the results of this task and the gap detection, it was determined that the optimal ISI for spatial tasks for B6 mice is 20 ms which is consist with other studies (Allen and Ison, 2010).

**Fig 3.**
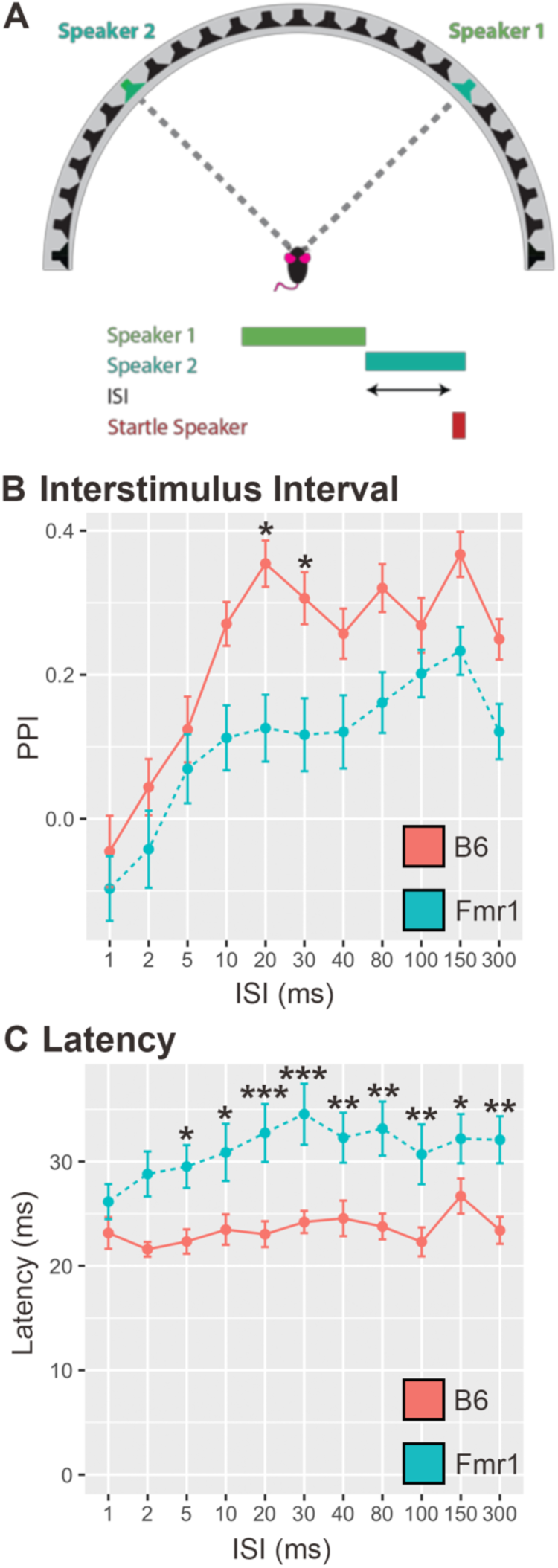
*Fmr1* mice have decreased PPI and increased startle latency in response to a 90*°* speaker swap compared to B6. 26 mice (12 B6, 14 *Fmr1*) were tested in a 90*°* angle speaker swap task with varying ISI (A). *Fmr1* mice (teal) have lower PPI at 20 ms and 30 ms ISIs than B6 mice (red; B). *Fmr1* mice had longer latencies to startle at all ISIs except 1 ms and 2 ms compared to B6 mice. Data are shown as mean ± SEM. * p<0.05, ** p<0.01, *** p<0.001.

### Minimum audible angle detection

To determine whether *Fmr1* mice have impairments in sound localization acuity, we measured and compared minimum audible angle in *Fmr1* and wildtype mice. The minimum audible angle is defined as the smallest change in speaker source location that the animals could just detect via the PPI metric above chance. In this task, the angle of the speaker swap across the midline was varied as the prepulse to the startle, with a constant ISI of 20ms as established by the previous experiments. 25 mice (11 B6 and 14 *Fmr1*) were tested with angle swaps (to the left and right of the animal) of 7.5, 15, 30, 45 and 90° across the midline with the animal oriented towards 0*°* (Figure 4A). Data were comparable for left to right and right to left directional swaps, therefore the data were pooled for both directions (Figure 4B, C). There was no main effect of genotype in this task (p=0.7255). *Fmr1* mice showed less PPI than B6 mice only at the 90*°* angle speaker swap, suggesting minimum audible was comparable in the two groups (Figure 4B). Both genotypes could not detect angles of 30° or less, as indicated by a PPI not significantly different than zero. These data suggest that both genotypes have poor minimum audible angle detection in this task. However, *Fmr1* mice had longer latencies to startle at all angles compared to B6 mice (Figure 4C) consistent with results in the previous varying ISI experiment.

**Fig 4.**
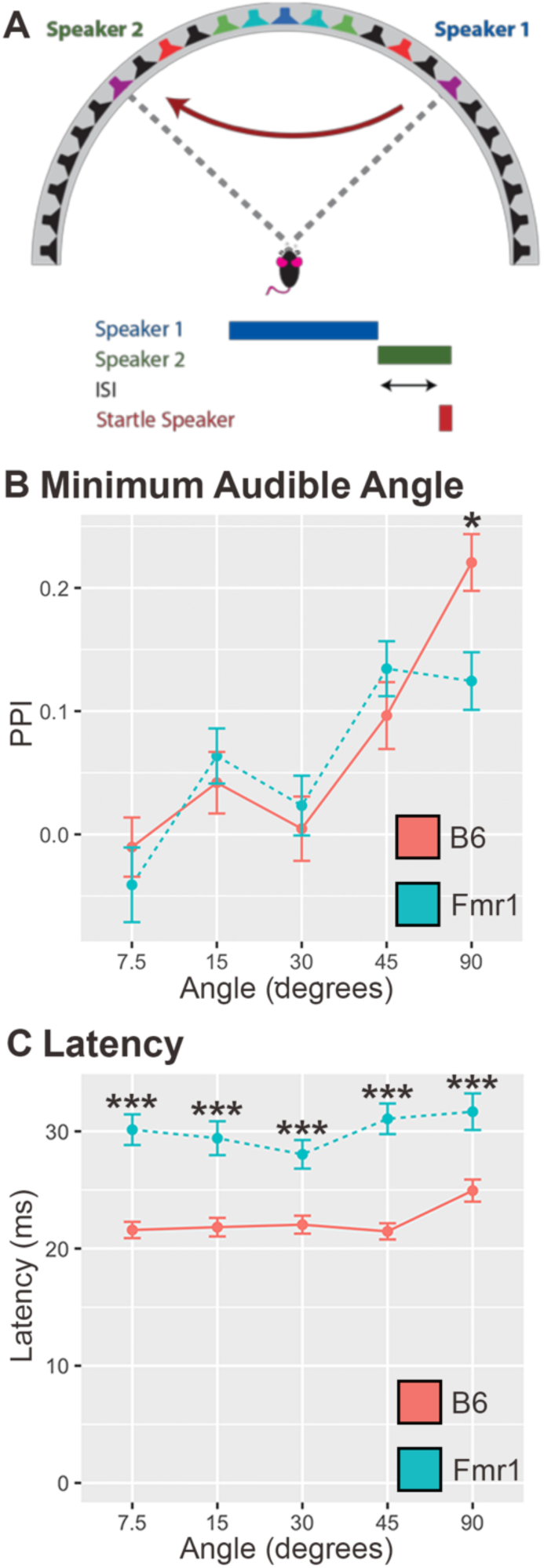
*Fmr1* mice had a reduction in PPI compared to B6 at 90*°* speaker swap and longer latencies to startle in all conditions. 25 mice (11 B6 and 14 *Fmr1*) were tested with varying angles of a speaker swap, both left and right directions across the midline, and constant 20 ms ISI (A, example leftward swap is shown). *Fmr1* mice (blue) show less PPI at 90*°* than B6 mice (red; B). *Fmr1* mice had longer latencies to startle at all angle conditions compared to B6 (C). Data are shown as mean ± SEM. * p<0.05, *** p<0.001.

### Auditory spatial release from masking-signal detection in noise

Listening to sounds in a complex auditory environment with competing sound sources (spatial release from masking) more naturally replicates real-world listening environments which people and mice with FXS experience and have difficulties with. We used the PPI task to replicate this experience and to determine whether *Fmr1* mice have impairments in spatial release from masking (SRM). First, we determined the signal attenuation required for animals to no longer be able to distinguish signals from the background. Spatial release from masking thresholds were measured in 32 animals (16 B6 and 16 Fmr1), which were placed in the chamber with a 70dB SPL masker sound presented from the 0° speaker and a beep centered around 4 kHz was played as the prepulse at varying attenuated levels from the adjacent speakers 7.5° to the left or right (Figure 5A). A beep was chosen because it is audible to the animal while not eliciting a social or emotional response (such as vocalization sounds etc. might). In this task, there was no main effect of genotype (p = 0.2786). At the loudest levels (lowest attenuation) *Fmr1* mice showed less PPI of their startle response compared to B6 mice (Figure 5B). Both genotypes did not detect attenuations of 24 dB and B6 animals did not detect attenuations of 27 dB, as indicated by a PPI not significantly different than zero. Similar to latencies in the gap detection task, B6 mice showed modulation of their latency based on condition, whereas *Fmr1* mice had consistent slower latencies across conditions (Figure 5C).

**Fig 5.**
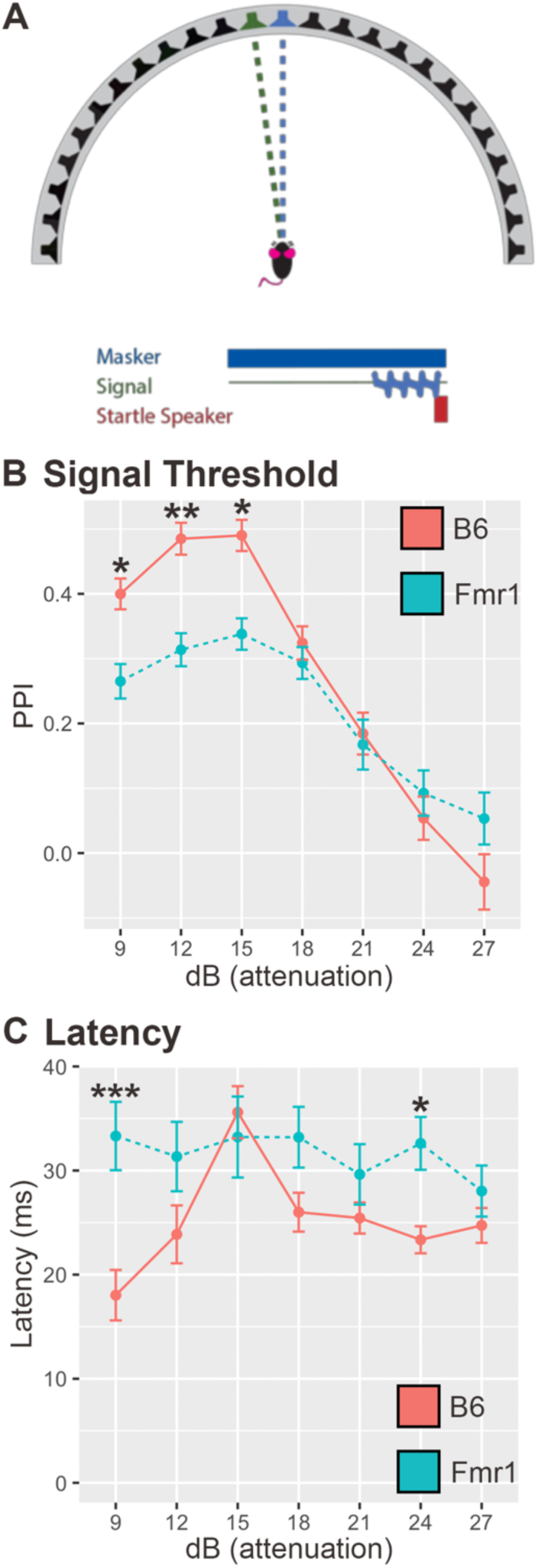
*Fmr1* mice showed less PPI of their startle, and longer latencies in some conditions compared to B6 mice. A signal noise was played at varying sound levels from a loudspeaker offset by 7.5° from a 70 dB SPL masker noise at 0° (A). *Fmr1* mice (teal) had reduced PPI of their startle at 9, 12, and 15 dBs attenuation compared to B6 mice (red; B). *Fmr1* mice also had increased latency to startle at 9 and 24 dBs attenuation compared to B6 (C). Data are shown as mean ± SEM. * p<0.05, ** p<0.01, *** p<0.001.

### Spatial Release from Masking Varying Angle

Wildtype animals showed reduced detection thresholds for sounds that originate some distance from a noise source, compared to those that are co-located. We wanted to determine if *Fmr1* mice had impairments in their angle detection as the signal approached the masker sound. In this task, 31 animals (15 B6 and 16 *Fmr1*) were presented with a signal (prepulse), at 2 attenuation levels (15 and 24 dB attenuation), at varying angles relative to a masker noise presented from the 0° speaker (Figure 6A). There was no main effect of genotype at either 15 dB attenuation (p = 0.1267) or 24 dB attenuation (p = 0.6325). Both *Fmr1* and B6 mice showed PPI greater than zero at all angles for the louder signal (15 dB attenuation), and only showed PPI greater than zero for the largest angle (90°) for the quieter signal (24 dB attenuation). Consistent with the speaker swap task, *Fmr1* mice only showed a difference in PPI relative to B6 mice at the largest angle (90°), and only at the louder (15 dB attenuation) level (Figure 6B, C). Similar to the above experiments, *Fmr1* mice had longer latencies to startle compared to B6 mice at several angles (7. 5 and 30° at 15 dB attenuation and 7.5, 22.5, 30, 45° at 24 dB attenuation Figure 6D, E). These data do not indicate substantial deficits in SRM; however, the longer latencies seen here, and in other experiments, suggests altered timing of startle responses in *Fmr1* mice compared to wildtype.

**Fig 6.**
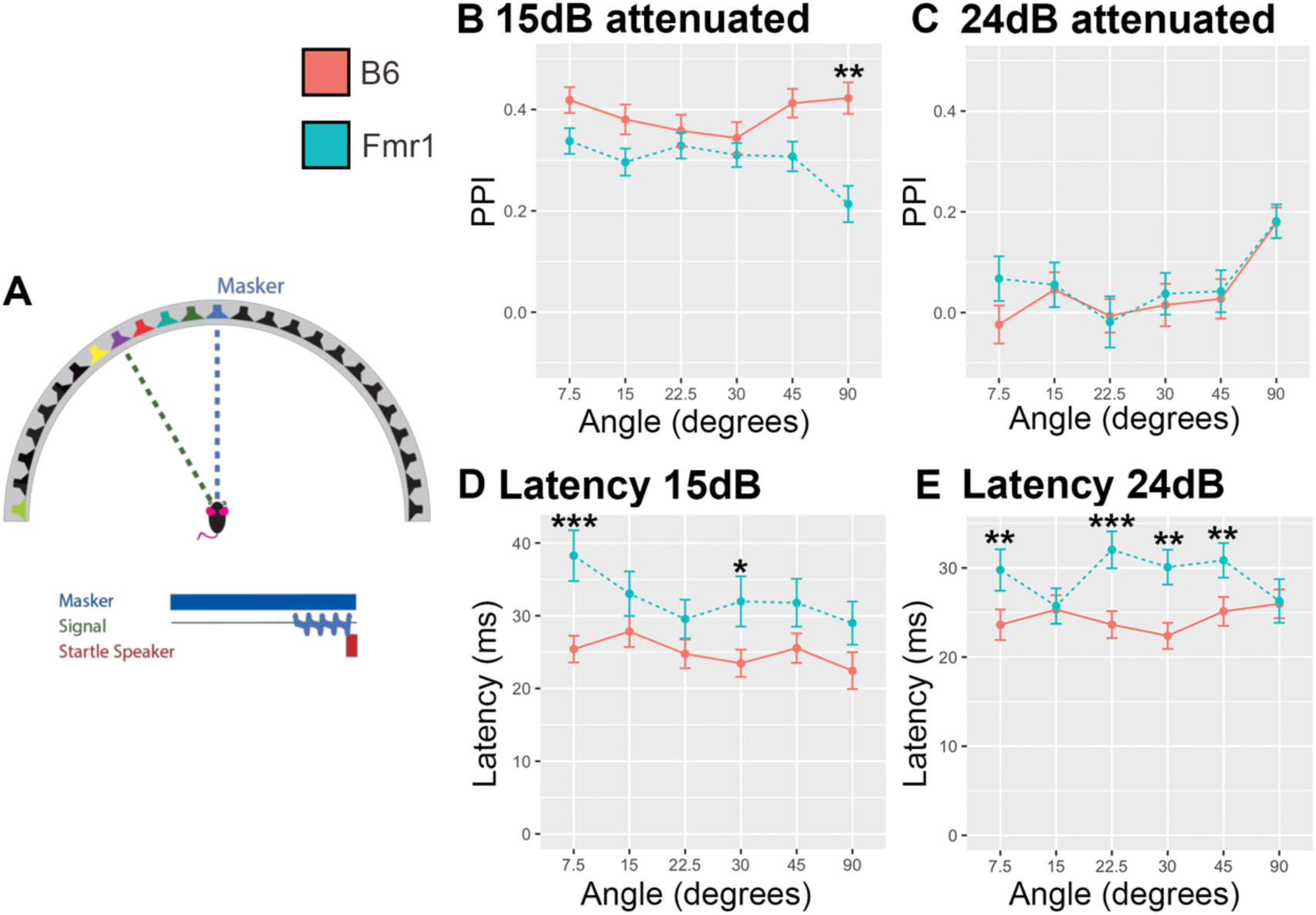
*Fmr1* mice had reduced PPI at 90° and 15 dB attenuation with increased latency to startle in several conditions compared to B6. The signal was varied at 2 levels of attenuation (15 and 24 dB) and varied at several angles from the 70 dB SPL masker at 0° to the left or right (A). *Fmr1* mice (blue) had decreased PPI of their startle response at 90° from the masker only at 15dB attenuation compared to B6 mice (red; B, C). *Fmr1* mice had increased latency to startle at 7.5 and 30° at 15 dB attenuation (D) and 7.5, 22.5, 30, 45, 90° at 24 dB attenuation (E) compared to B6 mice. Data are shown as mean ± SEM. * p<0.05, ** p<0.01, *** p<0.001.

### Habituation of the Startle Response

Previous studies have shown that *Fmr1* mice have impaired habituation (decreased startle response or PPI in later presentations of a replicate) during PPI tasks (Nielsen et al., 2002). Therefore, we used the replicate number per condition (ISI, dB, etc.) as the main effect variable per genotype. In contrast to previous studies, we found no habituation in either the B6 or *Fmr1* knockout mice in any of the experiments tested (startle threshold, gap detection, speaker swaps, or spatial release from masking)(Nielsen et al., 2002). This was indicated as no change in startle amplitude or PPI as a result of replicate, in particular the second replicate compared to the last replicate per condition block (p>0.05 in all tests).

## Discussion

This is the first detailed characterization of the binaural and spatial hearing ability of *Fmr1* mice. In addition, spatial hearing was tested using a non-invasive and reflexive PPI paradigm reducing potential complications of cognitive ability disparities between *Fmr1* and B6 mice. Surprisingly, *Fmr1* mice show spatial hearing ability comparable to wildtype mice, with only subtle differences noted in some prepulse conditions. In contrast, *Fmr1* mice had increased latency to startle in all experiments except when determining the startle threshold and overall acoustic startle response. These data suggest that perhaps *Fmr1* mice do not have a severe spatial hearing deficit but do show impairments in timing of responses. Lastly, we did not see any short-term habituation response within experiments in either the B6 or *Fmr1* mice.

### Startle threshold

Our results indicate that there is no difference in the overall acoustic startle response between *Fmr1* mice and B6 mice. However, when we assessed started force by accounting for the weight of the mice, we did see a difference at 100 and 110 dB SPL suggesting that *Fmr1* mice do startle with less force in at least some conditions compared to controls. Accounting for mass of an animal, at least in terms of startle threshold, could help interpretation of data across experimental methods and apparatuses (Grimsley et al., 2015). Previous studies of the acoustic startle response in *Fmr1* mice are equivocal, where some studies show an increase in the ASR ((Arsenault et al., 2016), at lower sound intensities (Nielsen et al., 2002)), while others show a decrease in the ASR (Baker et al., 2010; Chen and Toth, 2001; Frankland et al., 2004; Paylor et al., 2008; Spencer et al., 2006; Thomas et al., 2012; Veeraragavan et al., 2012; Yun et al., 2006; and Nielsen et al., 2002 at higher intensities), and others show no change in overall ASR (Ding et al., 2014) consistent with our study. The cause of these discrepancies is not clear; however, one possible explanation is the use of different mouse strains between previous studies. Different mouse strains can show varying ASR and PPI responses (Bullock et al., 1997). In particular, Chen and Toth, 2001 and Yun et al., 2006 used FVB background mice, whereas Frankland et al., 2004, Arsenault et al., 2016, Ding et al., 2014, Veeraragavan et al., 2012, Spencer et al., 2006, Paylor et al., 2008, Thomas et al., 2012 and Nielsen et al., 2002 used B6 background mice similar to our study. Nielsen et al., 2002 used an F1 generation cross of FVB and B6 knockouts which had an increase in ASR at low intensities and a decrease in ASR at higher intensities, and Baker et al., 2010 used an albino C57 strain.

Additional differences across reports include experimental set up and method of measuring the ASR. Each group used a different apparatus to assess ASR which could explain differing results, particularly when the magnitude of the effect is not dramatic. Furthermore, these previous studies did not account for weight of the animals by calculating startle force which, as we showed above, can significantly impact results (Grimsley et al., 2015).

Human studies of patients with FXS have also been conducted using the ASR. Neither Hessl et al., 2009 nor Frankland et al., 2004 found any differences in startle magnitude between patients with FXS and controls, which is consistent with our results when not corrected for weight. Lastly, studies have shown that the ASR is related directly to FMRP expression (Yun et al., 2006) and can be rescued with addition of the *Fmr1* gene (Paylor et al., 2008) suggesting that while the ASR phenotype is not that robust, some aspects of the ASR are directly related to loss of FMRP.

### Gap detection

Gap detection ability is thought to be directly related to E/I balance in the auditory system, particularly in the inferior colliculus (IC, Sturm et al., 2017), which is an important regulator of the ASR and PPI (Fendt et al., 2001; Koch, 1999). E/I imbalances have been found in *Fmr1* mice in areas of the auditory brainstem that contribute to sound localization processing, particularly the medial nucleus of the trapezoid body (MNTB, McCullagh et al., 2017; Rotschafer et al., 2015) and lateral superior olive (LSO, Garcia-Pino et al., 2017). These areas also contribute to the PPI and ASR pathways as they convey sound location information to higher areas such as the IC (Koch, 1999). Our data show that *Fmr1* mice show lower PPI magnitudes at short gap lengths (10 and 20ms) where the B6 mice are most robust at using the gap as a prepulse cue. This suggests that the *Fmr1* mice have impairments in their inhibition of the startle response leading to lower PPI compared to wildtype animals. In addition, our study examined latency to startle, and interestingly, *Fmr1* mice did not show the reduction in startle latency at gap ISIs longer than 20ms observed in wildtype mice. These data support the idea that imbalances in E/I at the level of the IC (and/or lower in the brainstem) lead to impairments in gap detection in *Fmr1* mice.

### Auditory spatial acuity

This is the first study to measure reflexively the minimum audible angle of *Fmr1* mice. We first examined if *Fmr1* mice inhibit their startle with an optimal ISI for a 90° speaker swap. Compared to B6 animals, *Fmr1* mice did not have as much PPI at any ISI indicating that they had trouble inhibiting the startle response even at such a large angle. In addition, when we kept the ISI at 20 ms and varied the angle, *Fmr1* mice again showed lower PPI values at 90° speaker swaps compared to B6 mice. However, both genotypes showed low PPI values for other speaker angles indicating that perhaps these mice did not have very good minimum audible angle perception in this task. Other studies have shown higher PPI values for mice at smaller angle swaps than we report, however differences in background strain (CBA/CaJ) and stimuli presented (wide-band noise vs. high-pass noise (>4 kHz, which according to the same study may still be low frequency for mouse)) may explain the differing results (Allen and Ison, 2010). In addition, consistent with other experiments reported here, the *Fmr1* mice responded with longer latencies to startle at almost all angle compared to B6. This indicates that not only do *Fmr1* mice have difficulties in inhibiting their startle response, they *Fmr1* mice may have longer processing speeds than B6 mice.

Operant conditioning paradigms can be more effective than PPI for assessing minimum audible angle (Behrens and Klump, 2016). However, in some disease models such as ours and other autism models, a reflexive task is preferred over one requiring training to remove any cognitive ability as a factor. In addition, Longenecker et al., 2016 showed that PPI was comparable to other measures (such as auditory brainstem responses) for assessment of auditory function. We show in our study that not only can we use PPI as a measure of minimum audible angle perception, we show impairments in minimum audible angle detection in *Fmr1* mice, albeit only at 90 degree angle swaps. These impairments may be even more pronounced in higher frequency noise or in more broadband noises as seen in the varying ISI experiment.

### Spatial Release from Masking

Spatial release from masking involves detecting a signal in a noisy background. Detecting a signal in competing noise may be the most applicable experiment to the real-world complex listening environment in which patients with FXS experience and have impairments. We show that even at loud signals compared to background, *Fmr1* mice respond with less PPI of their startle response than B6, suggesting their inhibition is less effective. In addition, when varying the location of the signal, at the louder sound, mice again had deficits at 90°, but this time off to the side of the animal, not a 90° swap in front of the animal (note, both measures test a 90° subtended angle across stimuli in the front hemifield). In contrast to the deficits seen with the 90° swap in front of the animal, deficits in this 90° off to the side may be due to the perception of the signal coming from slightly behind the animal. Depending on the location of their ear at the time of presentation (animals were constrained to face towards the 0° speaker but could move their heads slightly). Lastly, similar to the speaker swap experiments, *Fmr1* animals also had longer latencies under most conditions to respond to the startle speaker compared to B6, suggesting again not only impairments in detection, but also reaction ability/time.

Other studies have examined PPI while varying the intensity of a prepulse signal above an ambient noise level. While not exactly the same as the SRM task discussed here, in contrast to our results, most studies found that *Fmr1* mice had increased PPI compared to wildtype (Baker et al., 2010; Chen and Toth, 2001; Frankland et al., 2004; Nielsen et al., 2002; Paylor et al., 2008; Veeraragavan et al., 2012), however see also (Spencer et al., 2006; Thomas et al., 2012). These discrepancies could be due to the prepulse eliciting a startle response in these other studies, which would cause increased PPI during the actual startle, and in particular since these studies did not explore latency to startle, the prepulse startle response could be delayed coinciding with the startle-eliciting speaker. Lastly, most of the other studies do not discuss where the signal is coming from, which could impact the inhibition of the startle response in these animals and is likely different from our experiments. Interestingly, our results are consistent with data from patients with FXS who show reduced PPI under similar conditions (Frankland et al., 2004; Hessl et al., 2009). Consistency with human data implies that our assay may be a better measure of PPI that could apply to drug rescue experiments and be more applicable to the human FXS condition.

### Habituation

Often animals can habituate to the startle speaker, meaning that as the animal continues to be exposed to a loud sound stimulus, they will no longer startle as robustly as the earlier presentations of the stimulus. Habituation can also limit the length of experiments since animals may not respond as robustly after several hours of testing. Interestingly, we did not see any habituation to the startle in either *Fmr1* or B6 animals, as seen by a change in PPI or startle amplitude between early and later presentations of the same stimulus for any of the tasks presented. Other studies have examined habituation and shown that *Fmr1* mice do not habituate, though their results were not consistent between an F1 cross of genotypes and *Fmr1* mice on a B6 background suggesting that their results might be a result of background genotype (Nielsen et al., 2002). Additionally, impairments in habituation were present after ten presentations of the startle stimulus, most of our experiments had fewer than ten repetitions to avoid habituation to the startle response. Mice typically have less habituation than other animals, responding robustly and consistently to many stimulus presentations, and habituation can be extinguished with a few minutes rest between experiments (Valsamis and Schmid, 2011). Our data suggest also that mice can tolerate several hours of testing without a concern for habituation to the startle response, in particular in the B6 background strain tested here.

### Latency versus Startle

In contrast to PPI and ASR responses that differed between *Fmr1* and B6 mice, where we saw specific impairments under certain conditions, there was an overall trend for *Fmr1* mice to have increased latency to startle under a variety of conditions. This could be due to impairments in a different circuit that causes the response to the startle, i.e. when versus how much to startle in *Fmr1* mice. There has been one study which examined latency to react in patients with FXS after an acoustic startle, and they saw no differences between neurotypical controls and FXS (Roberts et al., 2013). This study however did not use PPI as a measure or look at EMG responses to the acoustic startle. None of the other studies examining PPI and ASR in *Fmr1* mice or humans examined the latency to respond making it difficult to know if our results are comparable. It would be interesting to explore the different pathways modulating the when and magnitude circuits to see if they are indeed modulated through different pathways and what implications that may have on the results discussed in this study on FXS.

### Conclusions

This is the first study to explore in depth the auditory deficits in an *Fmr1* mouse model of FXS using a reflexive PPI task. We saw minor differences in *Fmr1* mice compared to B6 mice in several measures of acoustic sound location (gap detection, minimum audible angle detection, and spatial release from masking) as measured by inhibition of the startle response. *Fmr1* mice also had increased latencies for almost all conditions compared to B6 mice suggested altered timing to acoustic cues. These experiments further show that, consistent with patient report and anatomical/physiological data, the auditory system is altered in a mouse model of FXS. In addition, this assay may be a useful tool for measuring drug efficacy on auditory impairments, in particular due to the similarity of responses between patient and mouse data.

## Materials and Methods

All experiments complied with all applicable laws, NIH guidelines, and were approved by the University of Colorado IACUC.

### Subjects

All experiments were conducted in either C57BL/6J background (wildtype) or hemizygous male and homozygous *Fmr1* knockout strain maintained on the background (commercially available through Jackson Laboratory, Bar Harbor, ME). Forty-six total mice were used, exact number of animals used per experiment are listed in the figure legend and corresponding results sections. Animals were genotyped regularly using Transnetyx (Cordova, TN). Mice used in these experiments were all adult animals and varied in age between 55 and 160 days old.

### Apparatus

Experimental conditions and apparatus are previously described in Greene et al., 2018 but are also briefly described here. All experiments were conducted in a double-walled sound attenuating chamber (IAC Bronx, NY) lined with acoustical foam to reduce echoes. The animal was snugly placed, in order to ensure the animal was forward-facing, in a custom-built acoustically transparent steel-wired cage attached to a polyvinyl chloride post anchored to a flexible polycarbonate platform with an accelerometer (Analog Devices ADXL335 Norwood, MA) to capture startle responses. All animals were tested in the dark using a closed-circuit infrared (IR) camera to monitor movement and proper orientation of the animal. When the animal was placed snugly into the chamber with the steel lightly compressed around its body, the animal maintained its forward-facing position and was unable to turn around. The cage with the animal was then always oriented towards the center loudspeaker. A diagram with the experimental apparatus is shown in (Greene et al., 2018). The chamber consisted of an array of 25 loud speakers (Morel MDT-20 Ness Ziona, Israel) placed horizontally in a 1m radius semicircular boom at 7.5° intervals from −90° (right) to +90° (left) in front of the animal from which pre-pulse stimuli were presented. Startle stimuli were presented from a Faital Pro HF102 compression driver (Faital S.p.A. San Donato Milanese, Italy) placed ~ 35cm (from the base of the platform) directly above the cage and amplified using an Alesis RA150 (Cumberland, RI).

Stimuli were generated and responses recorded from three Tucker-Davis Technologies (TDT Alachua, FL) RP 2.1 Real-Time processors using custom written MATLAB (Mathworks Natwick, MA) software. The startle stimuli were 20ms broad-band noise bursts generated by one of the RP2.1 processors and presented at 110dB SPL (unless otherwise stated such as in the startle threshold experiments). Carrier stimuli (CS) were broadband-noise generated by a second RP2.1 and presented continuously (unless otherwise noted) during testing. In the speaker swap experiment the broad-band noise was high-pass filtered with a 100^th^ order FIR filter. The CS was presented from one speaker at a time and had the ability to be switched by two sets of TDT PM2Relay power multiplexers controlled by the RP2.1s. Attenuation of signal was achieved using a TDT PA5 programmable attenuator. Vertical movement of the polycarbonate plate on which the accelerometer was mounted was detected as the voltage output, sampled at 1kHz by one of the RP2.1s, of the accelerometer. Startle response amplitude was calculated as the root mean square (RMS) of the accelerometer output in the first 100ms after the delivery of the startle stimuli.

### Experimental Conditions

Four types of experiment were conducted on most of the animals, startle threshold, gap detection, speaker swap, and spatial release from masking. The first repetition within a condition was excluded from analysis and all conditions were presented at least three times per experiment (this was only done with the startle threshold to determine if animals have a startle response) with most experiments containing five or more trials per condition. Most of the animals were tested once in each experiment; however, some mice were not subjected to acoustic startle threshold testing to reduce overall data collection time. The order of each of the four types of experiments was pseudo-randomized for each animal and each animal was only tested once per experiment (startle, gap, etc.). Total time of testing was around three hours per animal. The inter-trial interval (ITI time between trials) was uniformly distributed between 15-25 s in 1 s increments to prevent the animal from acclimating to the time of startle. For all experiments excluding the startle threshold, the startle stimulus was presented at 110dB SPL. Order of prepulse conditions were pseudo randomly presented for each experiment. For all experiments, latency to peak startle response within the first 100ms after the startle was also recorded.

### Experiment 1: Startle Threshold

Startle threshold was assessed by varying the intensity (for most animals between 60-120dB SPL in 10dB steps) of the startle eliciting stimulus, presented with the overhead startle speaker and recording their acoustic startle response (ASR) through the cage-mounted accelerometer. Startle responses were assessed in the presence of a 70dB SPL background noise played continuously from the speaker directly in front of the animal (0°). Presentation of these conditions were limited to three to five repetitions to ensure that the animal had a robust startle response while minimizing the duration of testing.

### Experiment 2: Gap Detection

The ability of animals to detect a short quiet period in a continuously noisy background was similarly assessed by presenting a broadband noise from the speaker directly in front of the animal (0°). A 20 ms gap in the noise (the pre-pulse) was introduced before the startle eliciting stimulus with interstimulus-intervals (ISI, time between the stimulus (gap) and startle)(Figure 2A) of 1, 2, 5, 10, 20, 40, 80, 160 and 240 ms from the onset of the gap. A subset of animals were only tested with 10, 20, 40, 80, 160 and 240 ms ISIs. Responses were assessed for ten repetitions of each ISI, presented pseudo randomly in a block-wise fashion. Two control condition trials, consisting of the continuous broadband noise with no gap preceding the startle-eliciting stimulus, were included in each repetition block.

### Experiment 3a: Varied inter-stimulus interval (ISI) with 90° Speaker Swap

The optimal ISI for speaker swap detection was assessed by swapping the source speaker (the prepulse) of a continuous broadband noise (70 dB SPL) 90° symmetrically across them midline (Figure 3A). The background noise was initially played from the speaker-45° (right) with respect to the animal and swapped with to the speaker +45° (left) of the animal some ISI prior to the startle-eliciting stimulus. Startle responses were assessed for five repetitions of 1, 2, 5, 10, 20, 30, 40, 80, 100, 150, and 300 ms ISIs, presented randomized in a blockwise fashion. Two control conditions, where no speaker swap occurred (i.e. the noise was continuously played from the initial speaker at −45°), were included in each repetition block.

### Experiment 3b: Speaker Swap

Minimum audible angle was similarly assessed using a speaker swap paradigm (Allen and Ison, 2010; Greene et al., 2018). The animal orientation was maintained at 0° (center) as described above to test responses to sounds swapped across the midline and assess minimum audible angle detection ability. The prepulse was a change in the source of a high-pass noise (cut off below 4kHz) between two matched speakers separated by 7.5°, 15°, 30°, 45°, and 90° symmetrically (except for 7.5°) across the midline, in both directions (left to right and right to left). ISI was set at 20 ms between presentation of the prepulse (speaker swap) and startle-eliciting stimulus. Startle responses were assessed for eight presentations of each condition (N = 10 since swap angle and direction co-vary), and one control condition (the high pass noise presented from the starting speaker, but no swap to the matched speaker) for each starting speaker (N=10), were presented randomized within repetition blocks.

### Experiment 4a: Threshold for Spatial Release from Masking

Spatial release from masking (SRM), i.e. the ability of mice to detect a signal in a continuous 70 dB SPL broadband masking noise, presented from the center speaker (0°), was assessed while varying the intensity of the “signal” speaker presented adjacent to the center speaker (7.5°, SRM threshold, Figure 5A). The ISI was set at 20 ms from the onset of the prepulse to the startle eliciting stimulus. The “signal” was a beep of 100 ms in duration centered around 4kHz by four octaves in frequency. The intensity of the signal was varied by decreasing the signal level by 9, 12, 15, 18, 21, 24, and 27 dB attenuation relative to the full scale, with a TDT PA5 programmable attenuator. Two control conditions (in which the masking noise was presented continuously with no “signal” prepulse presented). Were included in each of five randomized trial blocks.

### Experiment 4b: Speaker Swap Spatial Release from Masking

SRM was also assessed in a speaker swap task where instead of varying the intensity, the location of the signal speaker was varied at two levels of dB attenuation (15 or 24 dB attenuation Figure 6A). In this task, the same “signal” as above was presented at 15 or 24 dB attenuation, from speakers at 7.5°, 15°, 30°, 45°, and 90° (to the left or right) relative to center (0°) at a constant ISI of 20 ms. Two control conditions (in which the masking noise was presented continuously with no “signal” prepulse presented) were once again included in each randomized repetition block (of five).

### Data Analysis

The ASR was assessed as the RMS output of the accelerometer, amplified by 25dB in the 100ms following the startle eliciting stimulus presentation. The units of the ASR are given as units proportional to meters/second since the output of the accelerometer was not explicitly calibrated though it was held constant throughout data collection. The mean ASR was calculated for each animal, with the first presentation of each condition excluded to exclude initial adaptation to the startle. Most responses were quantified as prepulse inhibition (PPI), calculated as 1 minus the ratio of the mean prepulse ASR during each prepulse condition (ASR_p_) to the mean ASR during the corresponding control condition(s) (ASR_c_), recorded for each session: PPI = 1 – [ASR_P_/ASR_c_]. A PPI of 0 corresponds to an ASR_p_ equal to ASR_c_ suggesting no detection of the prepulse, whereas both positive PPI, indicating a reduction in ASR, and negative PPI, or an increase in ASR, suggest that the prepulse was detected and modified the animal’s startle response. Responses are presented as the mean of each of the populations tested (in this case B6 (wildtype) or *Fmr1* (knockout) mice) and error bars indicate the standard error of the mean (SEM). Figures were generated in R (R Core Team, 2013) using ggplot2 (Wickham, 2016). Data were analyzed using a mixed effects model to account for repeat observations within one animal (lme4 (Bates et al., 2014)) with genotype and conditions (dB SPL, ISI, angle etc.) as fixed effects, and animal as a random effect. Estimated marginal means (emmeans, (Lenth, 2019)) was then used to make pairwise comparisons between genotype and condition or replicate and condition. A tukey method for multiple comparisons was implemented for these contrasts using emmeans. A zero-intercept model was used to compare genotypes to a PPI value of zero (to determine detection of the sound prepulse). T-tests use Satterthwaite’s method for comparing the multiple levels of ISI across genotype (Kuznetsova et al., 2017). Animal weight was compared between the two genotypes using a two-tailed t-test. Where values are indicated as statistically significant, * indicated a p-value of <0.05, ** = p<0.01, and *** = p<0.0001. Figures were generated using Adobe Photoshop and Adobe Illustrator (Adobe, San Jose, CA).

## Acknowledgements

We would like to acknowledge Dr. John Peacock for his assistance in troubleshooting the various components of the behavioral set up. We would also like to acknowledge Dr. Peter McCullagh and Dr. Alex Kaizer for their assistance with the statistical models used in this paper.

## Abbreviations

ABR: auditory brainstem response
ASR: acoustic startle response
dB: decibel
E/I: excitation/inhibition balance
FMRP: Fragile X Mental Retardation Protein
FXS: Fragile X Syndrome
IC: inferior colliculus
ISI: inter-stimulus interval
ITI: intertrial interval
kHz: kilohertz
ko: knockout
LSO: lateral superior olive
MNTB: medial nucleus of the trapezoid body
PPI: prepulse inhibition
RMS: root mean square
SEM: standard error of the mean
SPL: sound pressure level
SRM: spatial release from masking

## Competing Interests

The authors state that they have no competing interests.

